# Point mutation in sensor triples zinc levels in *Sorghum bicolor*

**DOI:** 10.64898/2025.12.11.693441

**Authors:** Feixue Liao, Anko Blaakmeer, Søren Knudsen, Ana G. L. Assunção

## Abstract

Zinc (Zn) deficiency is widespread in agricultural soils and it is also a common nutritional problem in humans. We previously discovered a Zn Sensor Motif (ZSM) in the Arabidopsis F-bZIP transcription factors, bZIP19 and bZIP23, which impacts the concentration of Zn in Arabidopsis plants. Sorghum (*Sorghum bicolor (L*.*) Moench*) is one of the most important cereals crops worldwide and here, using a large-scale sorghum mutant library, we report the identification of variants with a ZSM point mutation. We show that a cysteine substitution in the ZSM of the sorghum bZIP19/23 homolog causes a three-fold increase in Zn levels in sorghum seeds with no visible developmental penalty. Our work provides the first evidence that modulating the F-bZIP ZSM is a promising strategy for Zn biofortification in crops. This discovery paves the way to generate Zn biofortified crops and alleviate the global problems of Zn deficient soils and Zn malnutrition.

## INTRODUCTION

Sorghum (*Sorghum bicolor (L*.*) Moench*) is one of the most important cereal crops in the world, being the dietary staple for over 500 million people. It is a versatile and fast-growing annual crop that thrives in warm and dry climates, including arid and semi-arid regions. It is widely cultivated in Asia, Africa and America, mostly for human consumption and animal forage and fodder, as well as bioethanol and biofuel production. The sorghum grain is gluten-free and a celiac-safe diet as well as its consumption and cultivation is expected to increase (Ananda *et al*. 2020; Khoddami *et al*. 2023). Sorghum is a primarily self-pollinating grass (Poaceae) with C4 photosynthesis, and a relatively small diploid genome (∼730Mb), which makes it an interesting crop for breeding varieties with improved nutrition value and climate resilience (Mason *et al*. 2024).

Deficiency of plant-available zinc (Zn) in soils is common globally, making Zn deficiency in crops a prevalent concern (Alloway 2008). The risk of Zn malnutrition affects approximately two billion people, particularly those living in developing regions with cereal-based diets. Deficiency of Zn impacts human growth and development, cognitive and immune functions, especially in children (Hussain *et al*. 2022). This global problem could be eased with breeding strategies to improve Zn use-efficiency and the Zn nutritional value (biofortification) of crops (White & Broadley 2009).

Plants maintain an adequate level of micronutrient Zn in their organs and cells through tightly regulated mechanisms of Zn homeostasis, which comprise Zn uptake from the soil, its transport, and distribution throughout the plant (Clemens 2001). In the model plant *Arabidopsis thaliana* (Arabidopsis), the transcription factors bZIP19 and bZIP23 were identified as the key regulators of Zn homeostasis and deficiency response. The double knockout mutant *bzip19 bzip23* were hypersensitive to Zn deficiency, while not showing a visible phenotype with sufficient Zn supply (Assunção *et al*. 2010). These transcription factors belong to the family of bZIP (basic region leucine zipper) transcription factors, which contain the characeteristic bZIP domain, comprising a basic region for DNA binding, and a leucine zipper for protein dimerization (Dröge-Laser *et al*. 2018). bZIP19 and bZIP23 belong to the F group within the bZIP family (F-bZIP), and regulate the response to Zn deficiency through transcriptional activation of Zn homeostasis related genes (**Fig 1a**). Target genes include members of the ZIP (ZRT/IRT-like Protein) family of Zn transporters, involved in cellular Zn uptake, and genes encoding the enzyme nicotianamine synthase (NAS), which catalyses the synthesis of nicotianamine (NA), a Zn ligand involved in the translocation and distribution of Zn in plants (Sinclair & Krämer 2012).

**Figure 1.**
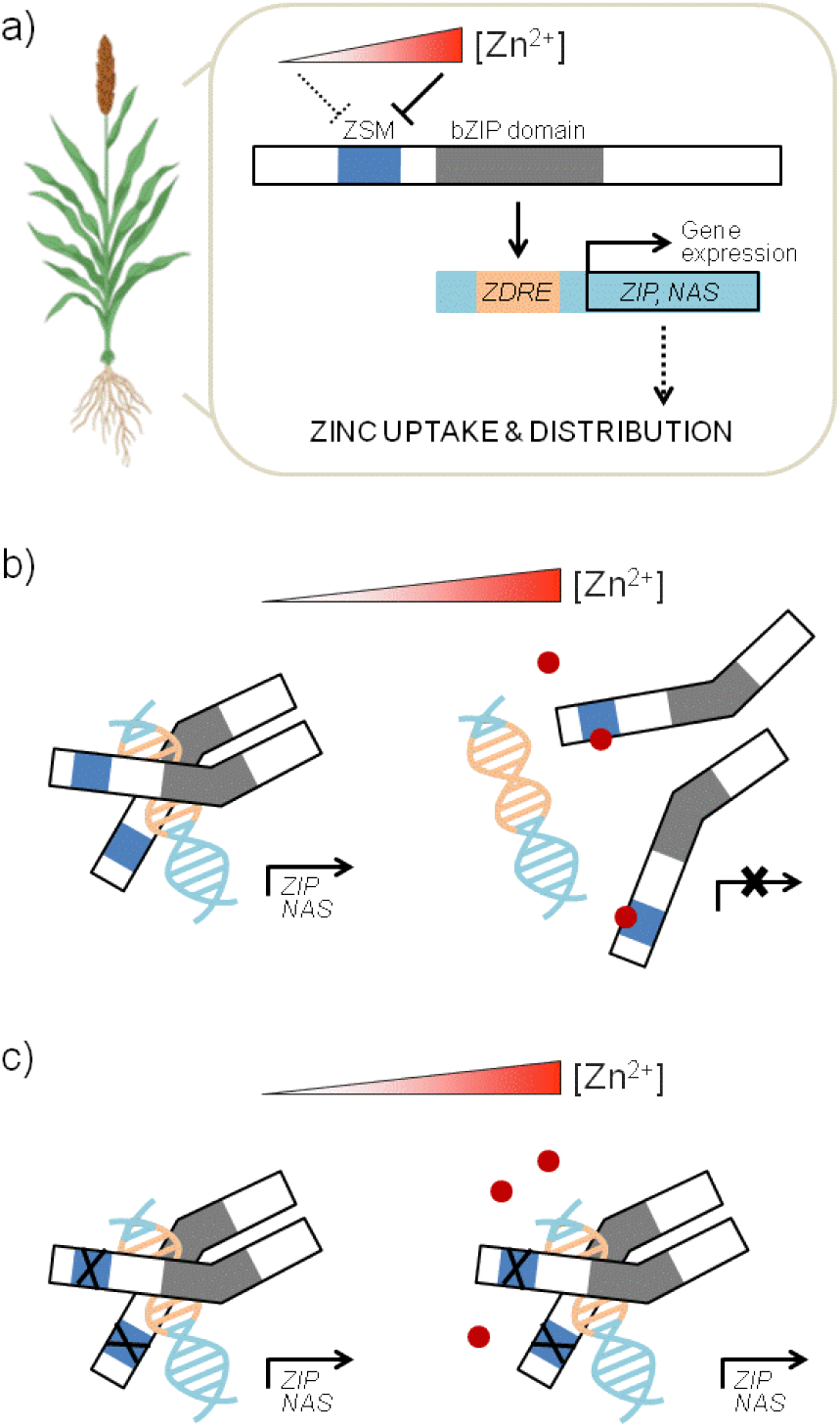
Zn deficiency response regulated by F-bZIP transcription factors. **a)** Simplified scheme depicting the transcriptional activation of target gene expression by the *Sorghum bicolor* F-bZIP transcription factor in response to Zn concentration status. ZSM and bZIP domain stand for Zinc Sensor Motif and basic region leucine zipper domain, respectively, in the F-bZIP protein. ZDRE is the Zinc Deficiency Response Element in the promoter of target genes. **b)** Simplified scheme depicting in more detail the regulation of F-bZIP activity by cellular Zn status. In short, under Zn deficiency, the transcription factor binds to the ZDRE in the promoter of their target genes, activating their transcription. These include *ZIP* and *NAS* genes, which encode Zn transporters and Zn ligands involved in Zn uptake and distribution. Under Zn sufficiency, cellular Zn ions bind to the Cys/His-rich ZSM in the F-bZIP protein halting the activity of the transcription factors. Zn^2+^ ions are depicted as red dots. **c**) Mutations in the ZSM (deletions or amino acid substitutions) lead to a Zn-insensitive constitutive transcriptional activation of the target genes. (adapted from Assunção 2022).

The bZIP19 and bZIP23 transcription factors also act as sensors of intracellular Zn status through binding of Zn ions to their Zn Sensor Motif (ZSM) (Lilay *et al*. 2021). The ZSM is rich in Cysteine (Cys) and Histidine (His) amino acids, which typically coordinate with Zn ions in Zn metaloproteins (Assunção *et al*. 2013). Under Zn sufficient conditions, cellular Zn ions bind the ZSM in nuclear localized bZIP19/23 halting the activation of target gene expression (**Fig 1b**). Interestingly, if the ZSM is deleted or mutated with amino acid substitutions, the Zn-regulated activity of the transcription factor is affected (**Fig 1c**). It becomes Zn-insensitive leading to a constitutive expression of the target genes and causing a significant increase of Zn concentration in Arabidopsis leaves and seeds (Lilay *et al*. 2021).

A phylogenetic analysis of F-bZIP homologs across land plants showed high level sequence conservation between F-bZIP proteins, and indicated conservation of the Zn deficiency response (Castro *et al*. 2017). F-bZIP proteins were already characterized as functional homologs of the Arabidopsis bZIP19 and bZIP23 in diverse land plants (Poaceae: rice (Lilay *et al*. 2020; Qing *et al*. 2024; Hu *et al*. 2024), wheat (Evens *et al*. 2017), barley (Nazri *et al*. 2017) and Fabaceae: *Medicago truncatula* (Liao *et al*. 2022)). The evolutionary conservation of the Zn deficiency response provides the opportunity for a translational approach from Arabidopsis to crops, targeting the ZSM of F-bZIP’s as a “molecular switch” for Zn biofortification (Assunção 2022).

For Zn biofortification to be effective, it is necessary to show that ZSM variants can substantially increase Zn accumulation in crops without compromising its performance. This demonstration has not yet been done. Sorghum is an ideal crop to evaluate the efficacy of ZSM variants in biofortification, because its genome has only one F-bZIP (*SB10G030250*) homolog to the Arabidopsis bZIP19/23 transcription factors. This *S. bicolor* F-bZIP member SB10G030250, now referred to as SbFbZIP1, was identified in phylogenetic analysis, and predicted to be the bZIP19/23 functional homolog conserving the Zn deficiency regulation function (Castro *et al*. 2017; Lilay *et al*. 2020). The sorghum SbFbZIP1 protein contains the bZIP domain, characteristic of bZIP transcription factors (Dröge-Laser *et al*. 2018). It also contains the ZSM at the protein N-terminus, characteristic of F-bZIP members (Castro *et al*. 2017) (**Fig 2a**).

**Figure 2.**
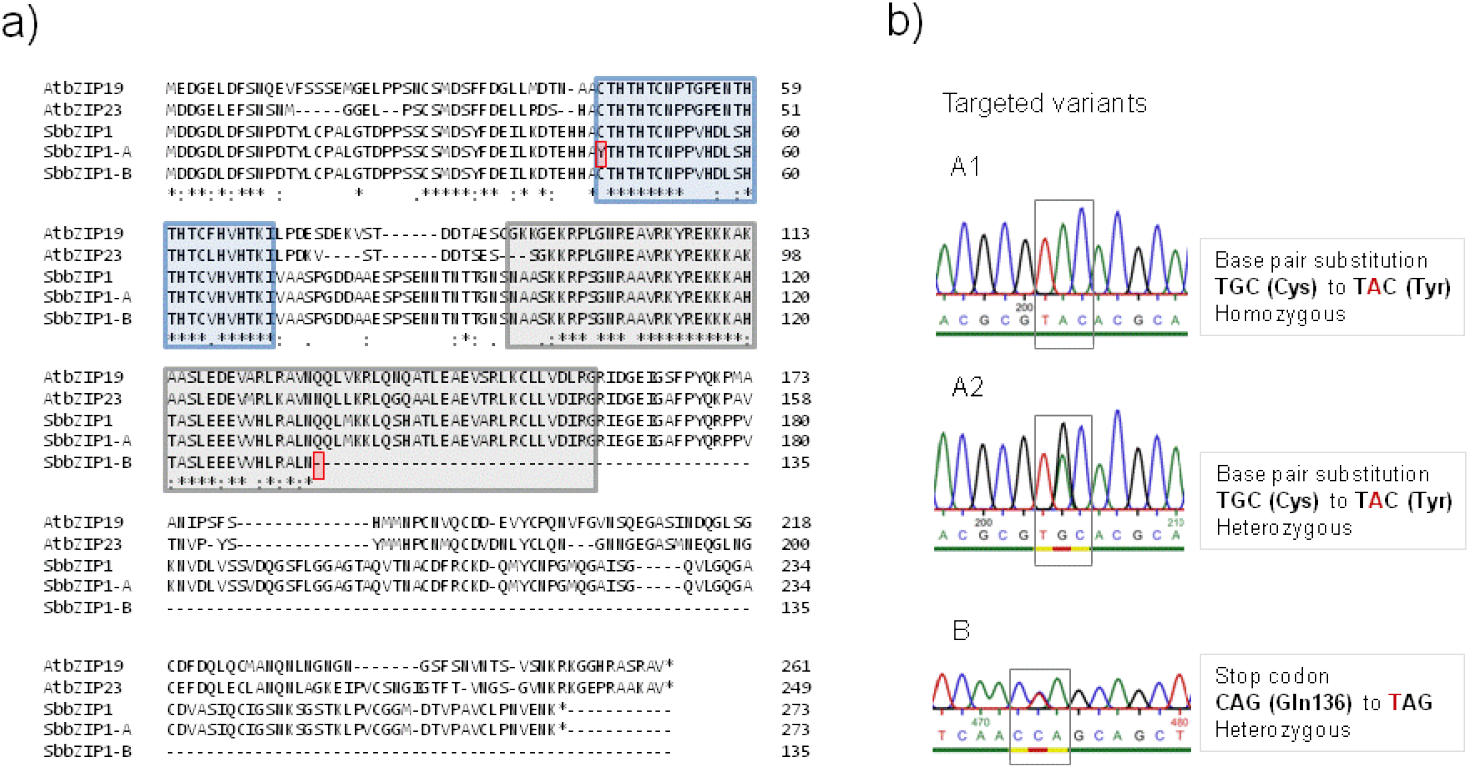
Sorghum SbFbZIP1 and the two point mutation genetic variants. **a**. Amino acid sequence alignment between Arabidopsis AtbZIP19 (AT4G35040) and AtbZIP23 (AT2G16770), sorghum SbFbZIP1 (SB10G030250), and sorghum variants SbFbZIP1-A and SbFbZIP1-B. The Zn Sensor Motif (ZSM) is shown in the blue box, and the amino acid substitution in the SbFbZIP1-A variant is highlighted in red. The bZIP domain is shown in the grey box, and the premature stop codon in SbFbZIP1-B variant is also highlighted in red. Asterisk * means that sequences have the exact same amino acid at that position. **b**. Sequence validation of selected M2 plants corresponding to variants A1, A2 and B. A snapshot of the sequencing chromatogram is shown corresponding to amino acid 45 (variants A1 and A2) and amino acid 136 (variant B).

Previously we showed, in Arabidopsis, that a single substitution of the first Cys residue of the ZSM leads to Zn-insensitive bZIP19 activity (Lilay *et al*. 2021). Here, we used a large-scale ethyl methanesulfonate (EMS)-induced sorghum mutant library (Mason *et al*. 2024), coupled with Fast Identification of Nucleotide variants by droplet DigITal PCR (FIND-IT) (Knudsen *et al*. 2022), to search for genetic variants in SbFbZIP1 either with a substitution of the first Cys of the ZSM, or with a premature stop codon (**Fig 2a**).

The EMS-induced mutagenesis targets primarily Guanine (G) and Cytosine (C) bases in the treated genome causing single base-pair substitutions from G to Adenine (A) and C to Thymine (T) (Sega 1984). The EMS treatment in the large FIND-IT populations is fine-tuned, on one hand, to induce single polymorphisms (SNPs) evenly distributed across genomes, and on the other hand, to be of low mutation rate to facilitate the recovery of mutant variants with minimal background genetic changes (Knudsen *et al*. 2022; Mason *et al*. 2024). It was estimated that all G/C bases in the genome are covered in this sorghum mutant library, meaning that all possible G/C mutations within the BTx623 genome are likely represented (Mason *et al*. 2025). This allowed screening for point mutations in *SbFbZIP1* that substitute the amino acid Cys (T**G**C) into Tyrosine (Tyr, T**A**C), and Glutamine (Gln, **C**AG) into a stop codon (**T**AG).

## RESULTS

### Sorghum library screening identifies SbFbZIP1 nucleotide variants

We identified two mutant plants for variant A, with the Cys45 to Tyr45 point mutation in the ZSM. These two M2 plants were found in the same mutant pool and therefore likely originate from the same M1 plant. We named them variants A1 and A2. For variant B, the Gln136 to stop codon point mutation, we identified one candidate. The M2 seeds of variants A1, A2 and B were germinated and allowed to grow. Sequencing analysis on M2 plants validated the expected single base-pair substitutions and, additionally, showed that variant A1 is homozygous whereas variants A2 and B are heterozygous for their respective single base-pair substitutions (**Fig 2b**).

We allowed M2 plants to self and produce M3 seeds, and we used M3 plants for trait evaluation analysis. Plants from M3 progeny were genotyped and the following were obtained: 5 variant A1 homozygous plants (Tyr/Tyr), 5 variant A2 homozygous plants (Tyr/Tyr), 2 wild-type plants from variant A2 (Cys/Cys), 2 variant A2 heterozygous plants (Cys/Tyr), 5 variant B homozygous plants (stop codon), and 5 non-mutagenized wild-type plants with the original BTx623 genome.

### A mutated ZSM controls Zn biofortification without visible penalties

To evaluate the Zn concentration trait in leaves and seeds, we grew the M3 plants on soil pots in a greenhouse for 12 months. We used ICP-OES to analyse Zn concentration in leaves (older and younger leaves separately) and tiller from 3-month-old plants, and in harvested mature seeds. Results showed that homozygous plants from variant B (B HM in **Fig 3**) did not differ in leaf and seed Zn concentration compared with wild-type (WT) plants (**Fig 3b**). This is expected because, although a premature stop codon likely represents a *sbfbzip1* knockout mutant, the F-bZIP-based Zn deficiency response is not activated in plants grown with sufficient Zn supply, as shown in Arabidopsis *bzip19/23* mutant (Lilay *et al*. 2019) (**Fig 1b**). Nonetheless, the 8-month-, and more noticeable, the 12-month-old plants from variant B showed more chlorosis than the wild-type (**Fig 3a, S1**) indicating the importance of an active F-bZIP regulator to cope with fluctuating Zn availability in long-term soil-grown plants, as was also shown for Arabidopsis (Huizinga *et al*. 2025).

**Figure 3.**
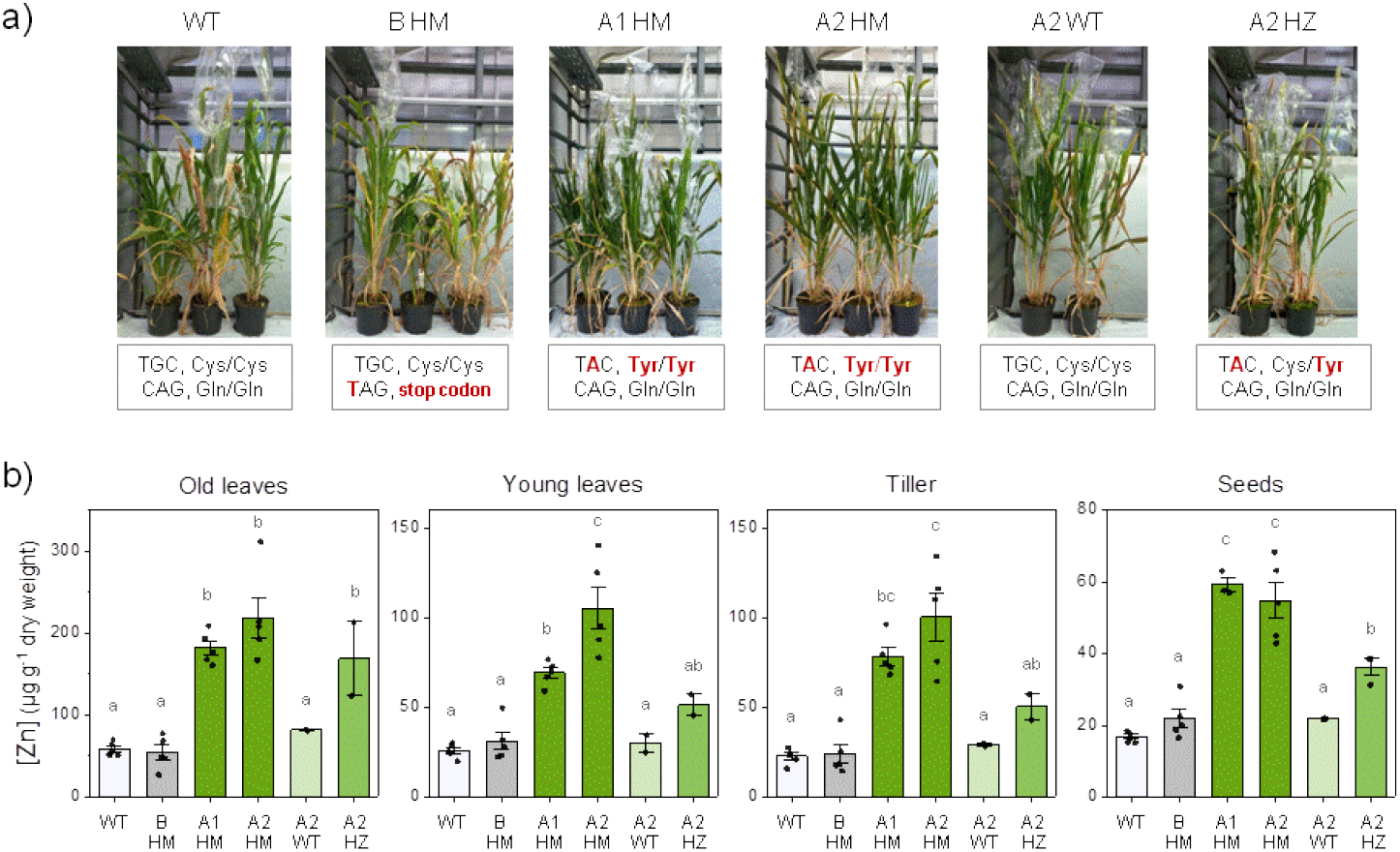
Analysis of sorghum variants. **a**. Images of 8-month-old sorghum plants, grown on soil in the greenhouse, depicting the genotypes BTx623 wild-type (WT) and variants B homozygous (B HM), A1 homozygous (A1 HM), A2 homozygous (A2 HM), A2 wild-type (A2 WT) and A2 heterozygous (A2 HZ). The corresponding base pair and amino acid substitution in each genotype are identified in a box below, with the point mutation identified in red. **b**. Average Zn concentration, measured with ICP-OES, in leaves (older leaves, younger leaves and tillers) harvested from 3-month-old plants, and in mature seeds. The individual replicas (n) are shown as data points; each datum corresponds to one leaf or tiller, or to 10 seeds, and each datum is from a different plant. For WT, BHM, A1HM, A2HM, n=5 plants, and for A2WT, A2HZ, n=2 plants, except for A1HM seeds with 3 samples from 1 plant. Data are mean ± s.e.m. Zn concentration in µg g^-1^ dry weight. Different letters indicate significant differences (P<0.05), determined using one-way analysis of variance followed by Tukey post hoc test.

In contrast, the homozygous A variants and the heterozygous A variant (A1 HM, A2 HM and A2 HZ in **Fig 3**) all had at least two-fold higher Zn concentrations in leaves, tiller and seeds than wild-type plants (**Fig 3b**). In the homozygous variants (A1 HM and A2 HM) this increase appears even higher, with plants showing at least a 3-fold higher Zn concentrations in leaves, tiller and seeds compared to the wild-type (**Fig 3b**). Analysis of other micronutrient concentrations (i.e., Fe, Cu, Mn) (**Fig S2**) and of the overall plant growth (**Fig 3a, S1**) indicate that this biofortification is specific to Zn, and that it is not associated with visible developmental penalties, as is also the case in Arabidopsis (Lilay *et al*. 2021; Huizinga *et al*. 2025). As expected, wild-type sorghum plants that segregated from the A2 variant (A2 WT in **Fig 3**) showed Zn content similar to that of non-mutated plants (**Fig 3b**). These results show that modulating the ZSM from SbFbZIP1, with a single Cys substitution, causes a large-effect Zn biofortification of sorghum plant and seeds. The results also confirm the expectations that the sorghum SbFbZIP1 is the functional homolog of Arabidopsis bZIP19/23 (Castro *et al*. 2017), and that ZSM modifications are gain-of-function mutations (Huizinga *et al*. 2025).

## DISCUSSION

This work provides the first proof-of-concept that modulating the ZSM from F-bZIP transcription factors with amino acid substitutions is an efficient “molecular switch” for Zn biofortification in crops, as previously hypothesized (Assunção 2022). (Qing *et al*. 2024) reported preliminary results showing increased Zn concentration in two rice lines with a gene-edited deletion of one or 19 ZSM amino acids from *OsbZIP4*, together with a gene-edited knockout of the other rice functional homolog of AtbZIP19/23, *OsbZIP50* (Lilay *et al*. 2020; Hu *et al*. 2024). However, these rice lines had severe plant developmental penalties. Here, in contrast, we show robust evidence of a Zn biofortification effect without apparent developmental penalties, resulting from a single nucleotide mutation at the sorghum F-bZIP ZSM. Our evidence is strengthen by the analysis of the segregating heterozygous A2 variant line, where the progeny plants segregating back to wild-type allele (Cys/Cys) lost the Zn biofortification trait (**Fig 3**).

Micronutrient content varies with environmental and growth conditions, but in general the concentration of Zn in sorghum seeds is around 20 mg kg^-1^ dw (Gaddameedi *et al*. 2022), in line with our results in BTx623 wild-type plants (**Fig 3b**). Currently, the target level for Zn biofortification in sorghum seeds is 32 mg kg^-1^ dw (HarvestPlus 2014; Guild & Stangoulis 2021), which is achieved with the Parbhani Shakti (14001), a released Zn biofortified sorghum variety (Gaddameedi *et al*. 2022). Our results with the homozygous A variants point to much higher Zn concentrations in sorghum seeds, in the range of 50-60 mg kg^-1^ dw (**Fig 3b**). Having found a strong Zn biofortification effect and no visible developmental penalties in the A variant, we anticipate a successful integration of the ZSM point mutation into elite plant germplasm to obtain sorghum plants biofortified with Zn and with increased use-efficiency in Zn deficient soils.

Given the magnitude of Zn deficiency problems in agricultural soils and human nutrition (HarvestPlus 2014), the importance of sorghum as a cereal crop, and the expected increase in its cultivation due to its resilience in a changing climate (Khoddami *et al*. 2023), the ability of developing Zn biofortified sorghum will impact nutrition security in regions with impoverished soils. Our findings unfold a direct link between a SNP and an important agricultural trait, highlighting the key contribution of plant functional biology and molecular breeding for finding novel strategies for a sustainable agriculture.

## Acknowledgments

This work was supported by the Independent Research Fund Denmark, Research Project 2 (DFF-FTP-9041-00182B) to F.L. and A.G.L.A.; Semper Ardens grant: “Crops for the future – Tackling the challenges of changing climates” from the Carlsberg Foundation (CF20-0352) to Birgitte Skadhauge at Carlsberg Research Laboratory. The authors wish to thank Morten Stephensen for support in the greenhouse facilities of the University of Copenhagen, and to thank Sandrine Chay for elemental quantifications at the “Service d’Analyses Multi-Elementaires “ (SAME) from the Institute for Plant Sciences from Montpellier, University of Montpellier/INRAE/CNRS/Institut Agro.

## Author contributions

F.L., A.B., S.K. and A.G.L.A. designed the experiments, F.L., A.B., and A.G.L.A performed experiments and analysed data, F.L. and A.G.L.A. wrote the manuscript revised by all authors.

## Competing interests

The authors declare no competing interests.

## SUPPLEMENTARY FIGURES

**Figure S1.**
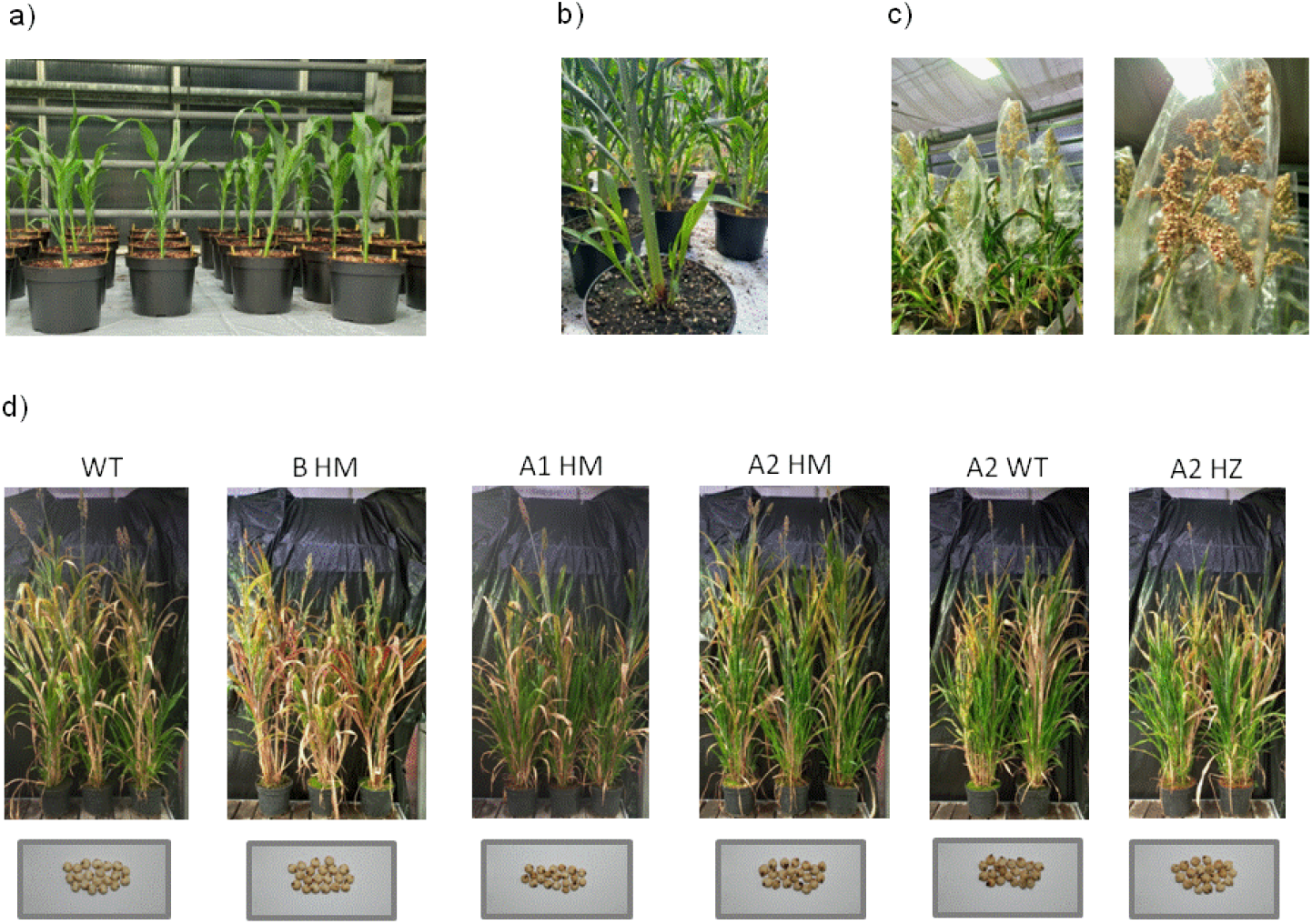
Sorghum plants at different growth stages in the greenhouse. **a**, Overview of 1-month-old plants. **b**, example of formed tillers in young plants. **c**, example of plants flowering and setting seed. **d**, images of 12-month-old sorghum plants depicting the genotypes, BTx623 wild-type (WT) and variants B homozygous (B HM), A1 homozygous (A1 HM), A2 homozygous (A2 HM), A2 wild-type (A2 WT) and A2 heterozygous (A2 HZ). Harvested seeds (n-20) from one plant from each genotype are shown in a box below.

**Figure S2.**
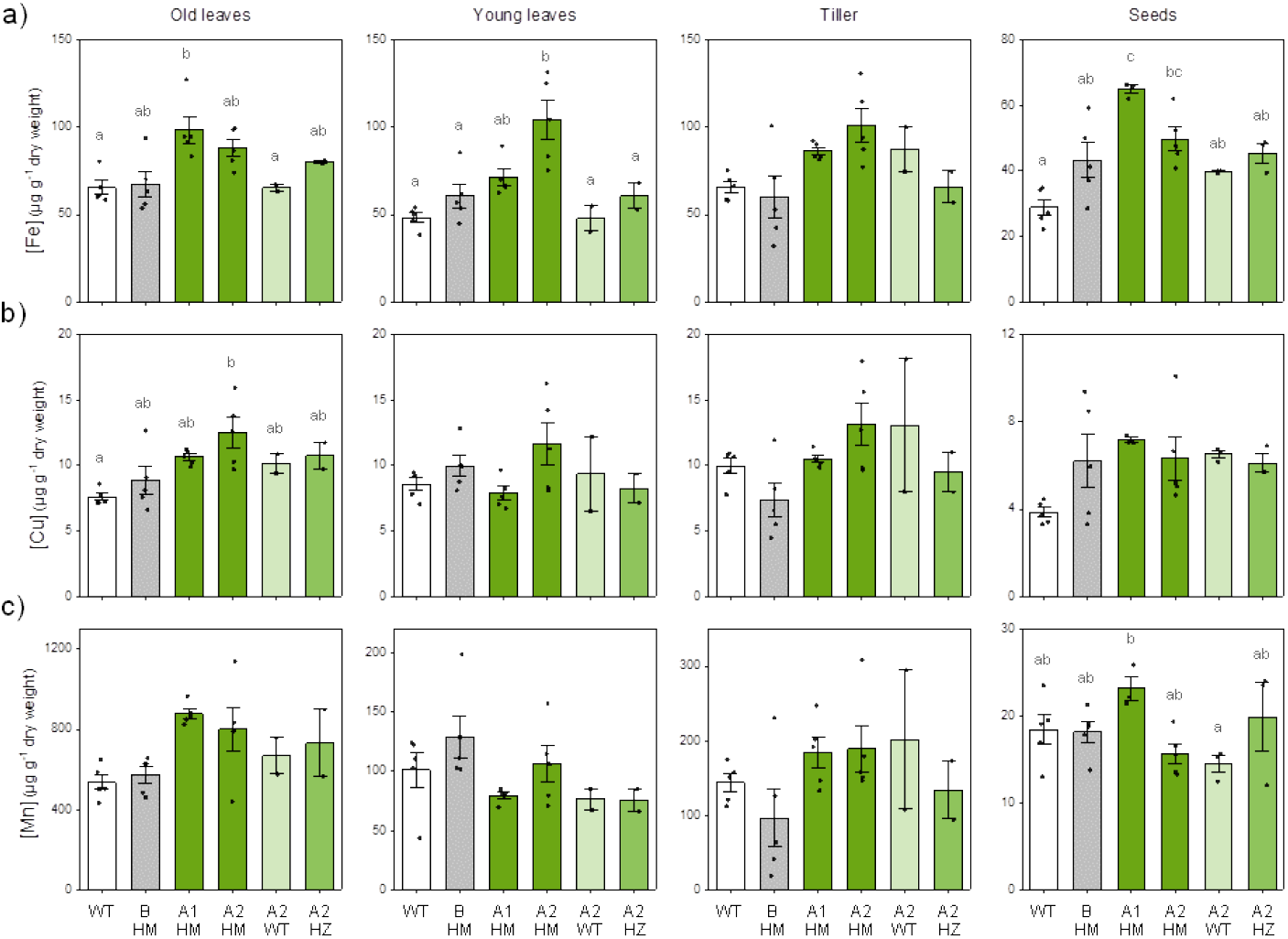
Tissue element analysis. Average concentration of (Fe) (**a**), cupper (Cu) (**b**) and manganese (Mn) (**c**) in leaves (older, younger and tillers) and seeds, using ICP-OES. The individual replicas (n) are shown as data points; each datum corresponds to one leaf or tiller, or to 10 seeds, and each datum is from a different plant. For WT, BHM, A1HM, A2HM, n=5 plants, and for A2WT, A2HZ, n=2 plants, except for A1HM seeds with 3 samples from 1 plant. Data are mean ± s.e.m. element concentration in µg g^-1^ dry weight. Different letters indicate significant differences (P<0.05), determined using one-way analysis of variance followed by Tukey post hoc test.

## METHODS

### Nucleotide substitution screening using FIND-IT technology

A large-scale sorghum M1 mutant library (150.000 plants) coupled with FIND-IT technology (Mason *et al*. 2024) was used to search for the nucleotide substitutions Cys45 (T**G**C) to Tyr (T**A**C), (variant A), and Gln136 (**C**AG) to a stop codon (**T**AG), (variant B), in the *SbFbZIP1 (SB10G030250)* gene. A detailed description of the FIND-IT protocol can be found in (Mason *et al*. 2024), but in summary; pooled DNA samples originating from seeds of 100 plants are screened until a positive pool is found (phase I). Seeds from the corresponding pool are drilled and flour is collected in batches of ten. DNA is isolated from the flour and screened with ddPCR (phase II). Seeds from the positive pools are grown and genotyped to identify the individual mutant plants (phase III). The primers used are (5’ to 3’): A_For (TCCATGGACAGCTACTTC), A_Rev (GTTGCAGGTGTGGGT), B_For (GAAGGCCCACACCG), B_Rev (GCTCTGGAGCTTCTTCA). The probes used are (5’ to 3’): A_HEX_WT (CACGCGT**G**CACGCA), A_FAM_MUT (ACGCGT**A**CACGCAC), B_HEX_WT (AGCTGCT**G**GTTGAGC), B_FAM_MUT (AGCTGCT**A**GTTGAGCG).

### Sequence validation

The identified M2 seeds were germinated, grown on soil and allowed to set M3 seeds. Leaf samples from the identified M2 plants were harvested for DNA extraction using the CTAB protocol followed by PCR amplification of *SbFbZIP1*. The primers used are (5’ to 3’): For (CTGATACCACTACAATTTTCC), Rev (CTAGAAGATAGACAGGAGTT). The amplified fragments were isolated and purified from agarose gel, and were sequenced. The analysis of the sequencing chromatogram allowed the genotyping and sequence validation of the M2 genetic variants, to confirm the mutations found with ddPCR. The same procedure was used to genotype M3 seedlings that were progeny of heterozygous M2 plants.

### Plant growth

Sorghum seeds were germinated on a Petri dish with wet filter paper in the dark at 28° C for 2-3 days. Germinated seedlings were planted on a soil with 60% peat and 40% vermiculite in 7,5 L pots, one seedling per pot. Plants grew for 12 months in a greenhouse under a 23 °C–18 °C light–dark cycle, with ventilation (windows) open at +2 °C, and with 16 hours of led light (blue 15%, green 35%, red 49% and far red 1%) at 250 µmol m^−2^ s^−1^, with the onset of curtains at 700 µmol m^−2^ s^−1^, and a relative humidity set at 65%. Plants were watered regularly with tap water, and once a week with soluble fertilizer (Pioner Basis Brown, pH 5.9, Ec. 2.2). From 3 month-old plants, tissue samples were harvested from each plant; one older and one younger leaf blade, cut from the collar, and one to two young tillers. Prior to flowering of the sorghum plants, the heads were bagged with micro-perforated pollination bags to ensure self-pollination. Upon maturation, seeds were harvested from each plant (**Fig S1**).

### Tissue element analysis

After harvest, samples (leaves and tillers) were placed separately in paper bags and dried in an oven at 60° C for 4 days, followed by grinded with zirconium bids in a high-speed shacking ball mill. The grinded samples and the seeds (10 seeds per plant) were digested in solution of HNO3 (65% [v/v]) and H2O2 (30% [v/v]) for a 3:1 ratio. Following an overnight incubation at room temperature, the samples were incubated at 85°C in HotBlock (Environmental Express) for 12 to 24 hours. Samples were then diluted 5 times by adding deionized Milli-Q water before the measurement. Elemental analysis was performed using Inductively Coupled Plasma Optical Emission Spectroscopy (ICP-OES 5800; Agilent Technologies) using calibration standard solutions provided by the manufacturer. The elemental concentration of each sample derives from 3 technical replicas. The ICP-OES is calibrated in wavelength each time the machine is switched on, using a 5 ppm calibration solution supplied by the manufacturer (ICP-OES&MP-AES wave cal 6610030100, Agilent Technologies) and performances tests are proceeded (optics, sensitivity and resolution). Quality control was performed with Certified Reference Material strawberry leaf powder (LGC7162), corresponded perfectly to the type of matrices usually handled. Atomic emission wavelengths are selected using the precision obtained from CRM measurements within +-10% of the theoretical value.

## Notes

### Competing Interest Statement

The authors have declared no competing interest.

